# FDAAA TrialsTracker: A live informatics tool to monitor compliance with FDA requirements to report clinical trial results

**DOI:** 10.1101/266452

**Authors:** Nicholas J. DeVito, Seb Bacon, Ben Goldacre

## Abstract

**Introduction:** Non-publication of clinical trials results is an ongoing issue. In 2016 the US government updated the results reporting requirements to ClinicalTrials.gov for trials covered under the FDA Amendments Act 2007. We set out to develop and deliver an online tool which publicly monitors compliance with these reporting requirements, facilitates open public audit, and promotes accountability.

**Methods:** We conducted a review of the relevant legislation to extract the requirements on reporting results. Specific areas of the statutes were operationalized in code based on the results of our policy review, publicly available data from ClinicalTrials.gov, and communications with ClinicalTrials.gov staff. We developed methods to identify trials required to report results, using publicly available registry data; to incorporate additional relevant information such as key dates and trial sponsors; and to determine when each trial became due. This data was then used to construct a live tracking website.

**Results:** There were a number of administrative and technical hurdles to successful operationalization of our tracker. Decisions and assumptions related to overcoming these issues are detailed along with clarifications directly from ClinicalTrials.gov. The FDAAA TrialsTracker was successfully launched in February 2018 and provides users with an overview of results reporting compliance.

**Discussion:** Clinical trials continue to go unreported despite numerous guidelines, commitments, and legal frameworks intended to address this issue. In the absence of formal sanctions from the FDA and others, we argue tools such as ours - providing live data on trial reporting - can improve accountability and performance. In addition, our service helps sponsors identify their own individual trials that have not yet reported results: we therefore offer positive practical support for sponsors who wish to ensure that all their completed trials have reported.

## Introduction

The results of clinical trials are used to inform treatment choices. Complete reporting of all clinical trial results is widely recognized as a clinical and ethical imperative.^1,2^ However it has long been documented that trial results are left undisclosed^3^ and the most current systematic review of publication bias cohort studies shows that only half of all completed trials on registries report results^4^, consistent with earlier work.^5^

There is now a growing movement towards legislation requiring results to be reported online, within 12 months of completion, on both EU^6,7^ and US^8,9^ registries. In January 2018 the first trials to be covered by updated US trial reporting regulations, under the Food and Drug Administration (FDA) Amendments Act of 2007 (FDAAA 2007), became due to report results. This is a potentially important legal landmark, against a background of slow and incomplete progress on trials transparency.^10^

As there is currently no official accounting of trials covered under the FDAAA 2007 and their reporting status, we therefore set out to develop and deliver an online tool which publicly monitors compliance with these new reporting requirements, facilitates open public audit, and promotes accountability.^11^ While it is important that *all* trials are reported, we set out to identify, document, and faithfully implement all the exceptions of FDAAA 2007, to ensure we only identify trials covered by this legislation as breaching to the extent possible using publicly available information available from ClinicalTrials.gov.

## Methods

Our specific objectives were: to review the legislation; create processes to automate the download and management of ClinicalTrials.gov data; to develop a method to identify due trials in the data; and to deliver an online interactive web platform presenting FDAAA 2007 compliance data to users.

### Policy Review

A policy review was conducted to ascertain the relevant reporting requirements of FDAAA 2007 and Final Rule 42 CFR Part 11 of 2016 (Final Rule).^8,9^ Additional materials, related to interpretation and implementation of these statutes, available directly from ClinicalTrials.gov, were also reviewed.^12–15^ Any further questions on the reporting requirements and their implementation on ClinicalTrials.gov were referred to the ClinicalTrials.gov team through their official “Customer Support” channel.^16^ All communications with ClinicalTrials.gov were archived and are available as Appendix 1.

### Obtaining the Data

ClinicalTrials.gov provides an updated XML record of the ClinicalTrials.gov database each day.^17^ This data was downloaded and used to create a queryable database on Google’s Big-Query platform for prototyping. The full ClinicalTrials.gov dataset from each download is processed and archived.

### Interpretation and Implementation

Prototyping for data extraction and applicable trial identification was conducted using Big-Query (Standard SQL). Specific areas of the statutes were operationalized in code based on the results of our policy review and the publicly available data elements on ClinicalTrials.gov. We developed methods to identify trials required to report results using publicly accessible data; to collect additional relevant information; and to determine when trials became due, using key trial dates. Once prototyping was complete, all data processing procedures and trial identification logic was converted to Python 3 code. When faced with any issues of data availability or interpretation, every effort was made to conservatively assess whether trials are covered, and when they were due in order to limit false positives.

### Web Tool

Our dataset and code was used to create a regularly updated website (fdaaa.TrialsTracker. net) to display all Applicable Clinical Trials (ACTs) and probable Applicable Clinical Trials (pACTs); track when they become due; show whether they have reported results in accordance with the law; give performance statistics for each individual trial sponsor; and calculate potential fines that could have been levied by the FDA against sponsors.

### Data and Code Sharing

All underlying code related to data extraction and website development is made freely available for review and re-use under the MIT open source license via public GitHub repositories.^18,19^

## Results

### Policy Review

#### Background to FDAAA 2007 and Final Rule

The FDAAA 2007 required that certain trials share their results on ClinicalTrials.gov.^8^ While the global ethical standard is that all trials should report results^1,2^, this legislation provides numerous reporting exceptions that exempt certain clinical trials from their obligation to report. The initial reporting requirements of FDAAA were vague and left some details open to interpretation regarding who was required to report and when.^20–22^ It was not until 2016, with the publication of the Final Rule^9^, that these requirements were further clarified and expanded: specifically, they state that all trials of both approved and unapproved products, meeting various clearly specified criteria, are required to report results within one year of their completion date. The Final Rule also created more straightforward ways to determine which trials are classed as “applicable” and hence due to report, including specifying new criteria for ACTs.^23^ These new standards came into effect on January 18, 2017.

#### Identifying ACTs & pACTs

In order to identify which trials are required to report results, it was necessary to determine which trials are ACTs or pACTs. An ACT is any “applicable trial” which began on or after the effective date of the Final Rule; an applicable trial is determined using the criteria in Table 1. The term “probable ACT” (pACT) is officially used to denote a trial which began prior to, but ends on or after, the effective date of the Final Rule and based on the available information is likely covered by the law (again as per Table 1). Because certain data elements required to identify ACTs were either not available or not required prior to the implementation of the Final Rule, pACTs are identified in the Final Rule using a separate methodology from ACTs. These criteria are also officially documented in the ClinicalTrials.gov Protocol Registration and Results System (PRS) User’s Guide which notes that “records that meet the [ACT/pACT] condition...will be flagged for FDAAA or 42 CFR Part 11 issues.”^14^

**Table 1:**
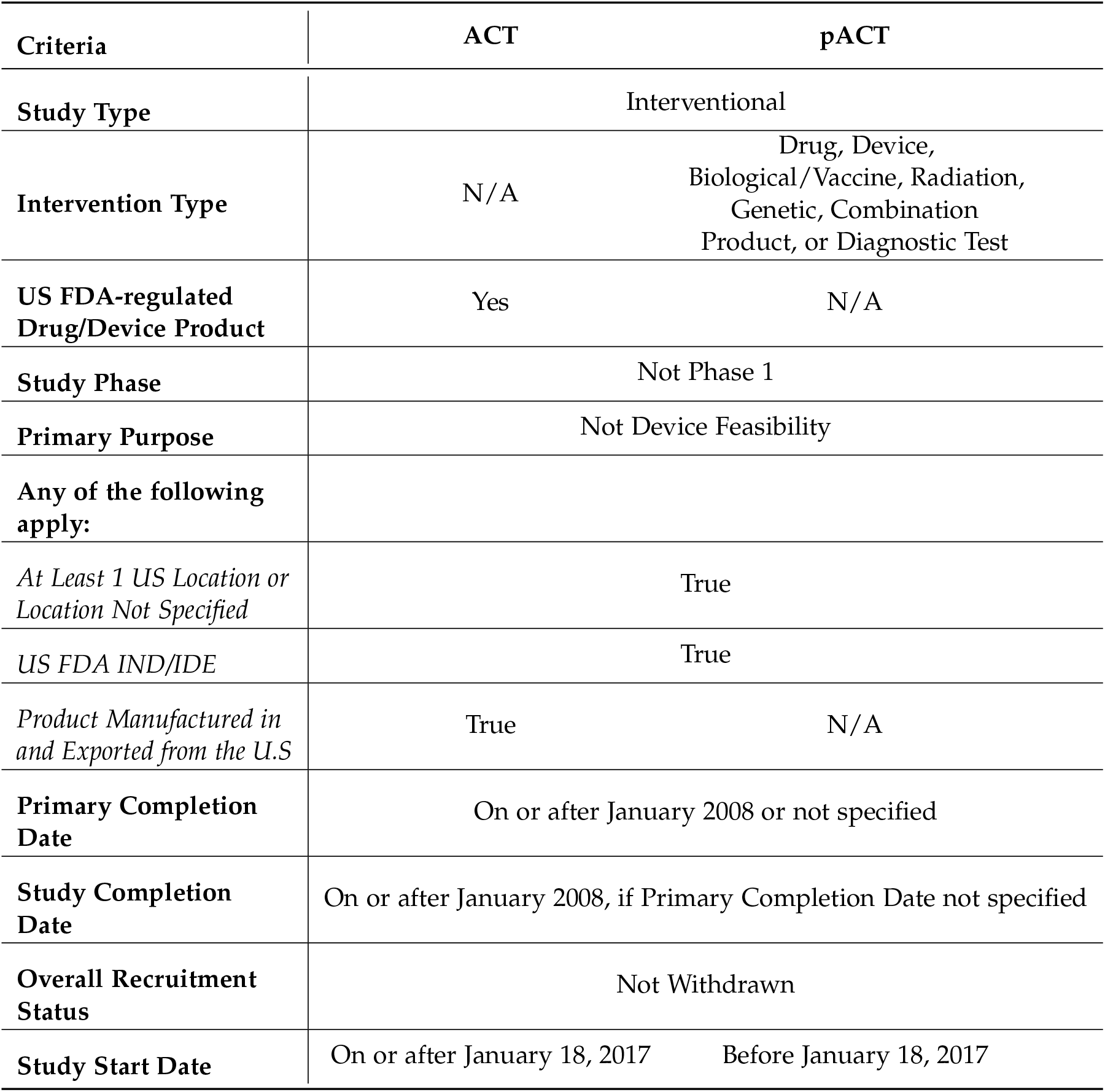
ClinicalTrials.gov Criteria for ACTs and pACTs^14^

| Criteria | ACT | pACT |
| --- | --- | --- |
| **Study Type** | Interventional |  |
| **Intervention Type** | N/A | Drug, Device,<br>Biological/Vaccine, Radiation,<br>Genetic, Combination<br>Product, or Diagnostic Test |
| **US FDA-regulated**<br>**Drug/Device Product** | Yes | N/A |
| **Study Phase** | Not Phase 1 |  |
| **Primary Purpose** | Not Device Feasibility |  |
| **Any of the following**<br>**apply:** |  |  |
| *At Least 1 US Location or*<br>*Location Not Specified* | True |  |
| *US FDA IND/IDE* | True |  |
| *Product Manufactured in*<br>*and Exported from the U.S* | True | N/A |
| **Primary Completion**<br>**Date** | On or after January 2008 or not specified |  |
| **Study Completion**<br>**Date** | On or after January 2008, if Primary Completion Date not specified |  |
| **Overall Recruitment**<br>**Status** | Not Withdrawn |  |
| **Study Start Date** | On or after January 18, 2017 | Before January 18, 2017 |

An interesting barrier is presented by the fact that, although ClinicalTrials.gov and the FDA hold data on which trials are ACTs or pACTs, they do not share this information publicly. However, public documentation identifies all data elements used to determine ACT and pACT status.^9,12,14^ The PRS criteria are accompanied by the following caveat:

> The PRS identifies records that appear to be probable ACTs (pACTs) or ACTs based solely on information submitted for the data elements listed below. These records should be carefully reviewed, but the records identified with FDAAA or 42 CFR Part 11 issues may not be comprehensive (it may include records for trials that are not ACTs or exclude records for trials that are ACTs).

Operationalising these criteria was itself complicated by the fact that Investigational New Drug (IND) and Investigational Device Exemption (IDE) status is a required element to fully identify ACTs and pACTs, but is not available in the public dataset for any trial.^13^ However, this can be worked around: outreach to ClinicalTrials.gov support confirmed that for ACTs the “FDA Regulated Drug/Device” criteria cannot be entered as “Yes” during trial registration (or subsequent updates) unless the trial either involves a US location, is conducted under an IND/IDE, or the product is manufactured in and exported from the US (Appendix 1). We therefore only included “FDA Regulated Drug/Device” status in our ACT logic in lieu of these additional criteria. This is in line with language from the final rule that states: “Promulgation of the final rule and implementation of several new data elements (e.g., Studies an FDA-regulated Drug [or Device]), enables the Agency to be better able to identify applicable clinical trials more accurately in the PRS and on the public Web site.”^9^

“FDA Regulated Drug” and “FDA Regulated Device” are new data elements only available since the implementation of the Final Rule and only required for ACTs. pACTs may choose to update and include these fields but it is not required. Originally, our pACT criteria required a trial to have a US location, and one of a number of specified intervention types, in alignment with the pACT criteria in the PRS UserâĂŹs Guide. Following outreach to ClinicalTrials.gov in January 2019 (Appendix 1), we felt confident that when the “FDA Regulated” fields are present, we could disregard these pACT-only fields in favour of the ACT criteria. While ClinicalTrials.gov would not directly confirm that the same assumptions could be made about the “FDA Regulated” fields for both ACTs and pACTs, they pointed us towards language in the ClinicalTrials.gov FAQ for determining ACT coverage which we believe supports this decision:^24^

> Beyond their primary purpose, the ACT Checklist and Elaboration may also be useful to assist in evaluating whether a clinical trial or study that was initiated before January 18, 2017, and which is not subject to the final rule requirements, is an ACT under section 402(j) of the Public Health Service Act.

When these fields are not present, we default to the original pACT criteria, which includes intervention type and the US location requirement. This approach will exclude some pACTs that provide no US location, or no locations at all, and have an IND/IDE that is not flagged in ClinicalTrials.gov data: this is conservative, because some trials giving no location may in reality be conducted in the US, but not be identifiable as such, because the sponsors have entered poor quality data onto the register. This change went into effect in March 2019 and added 300 new trials to the Tracker.

Prior to the implementation of this new post-2017 FDA Regulation field, ClinicalTrials.gov contained an older field named: “is_fda_regulated”. This was deleted from ClinicalTrials.gov on January 11, 2017.^17^ While the old field and the new fields functionally convey the same information^9^, no information was preserved from the deprecated field in the current ClinicalTrials.gov dataset; we regard this as a sub-optimal approach to data stewardship for a public resource. Without any data for the old or new fields concerning FDA regulation there is no way to exclude a pACT from reporting requirements due to their FDA regulation status. An archived copy of the ClinicalTrials.gov database from January 5, 2017 is available via the Clinical Trials Transformation Initiative.^25^ From this dataset we were able to extract the “is_fda_regulated” data for all trials as it existed immediately prior to the removal of the field. In a manual review of pACTs which had previously used the “is_fda_regulated” field, it was determined that utilizing this data would provide an additional useful exclusion criterion for pACTs. This field did not, however, appear to be entirely accurate as some of the trials reviewed appeared as if they should be required to report. However, to maintain our conservative approach, all trials identified as not being FDA regulated in the January 5, 2017 dataset will be excluded from the tracker unless more recent information on FDA-regulation status becomes available. Trials that are identified as being “FDA regulated” by the archived field are not, however, automatically included unless they meet all other pACT criteria including an explicit US location.

The criteria in Table 1 also identify post-2008 completion dates as required criteria for both pACTs and ACTs. All pACTs and ACTs relevant to our tracker will have completion dates on or after January 18, 2017 so this criteria was unnecessary for our purposes. While the official ACT/pACT criteria also includes trials with no completion date specified, it is impossible to track Final Rule compliance without a completion date to anchor the 12 month reporting window, and therefore these trials cannot be included in our tracker. Table 2 shows our final logic for determining ACTs and pACTs based on the public data.

**Table 2:** ACT and pACT Logic

| Category | Logic |
| --- | --- |
| **ACT Logic**<br>(Trials started on or After 18 Jan 2017)<sup>a,b</sup> | Study Type is *Interventional*<br>**AND**<br>FDA Regulated Drug **OR** Device is Yes<br>**AND**<br>Phase is 1/2, 2, 2/3, 3, 4 or N/A<br>**AND**<br>Primary Purpose is **NOT** *Device Feasibility*<br>**AND**<br>Study Status is **NOT** *Withdrawn* |
| **pACT Logic**<br>(Trials started prior to, but completed on or after, 18 Jan 2017)<sup>a,b</sup> | Study Type is *Interventional*<br>**AND**<br>Phase is 1/2, 2, 2/3, 3, 4 or N/A<br>**AND**<br>Primary Purpose is not *Device Feasibility*<br>**AND**<br>Study Status is not *Withdrawn*<br>**AND**<br>IF FDA Regulated Drug **OR** Device Field is Available:<br>FDA Regulated Drug **OR** Device is Yes<br>IF FDA Regulated Drug **OR** Device Field is **NOT** Available:<br>Intervention Type is *Biological OR Drug OR Device OR Genetic OR Radiation OR Combination Product OR Diagnostic Test*<br>**AND**<br>Study Location includes *United States* or *US Territories*<sup>c</sup><br>**AND**<br>"Is FDA Regulated" is *True OR Null*<sup>d</sup> |

<sup>a</sup> For all date values, when only a Month/Year were given, dates were defaulted to the last day of the given month (e.g. January 2017 = January 31, 2017).
<sup>b</sup> "Completion Date" field was used when "Primary Completion Date" was not available.
<sup>c</sup> "US Territories" include American Samoa, Guam, Northern Mariana Island, Puerto Rico and the U.S. Virgin Islands.
<sup>d</sup> "Is FDA Regulated" field available via archived ClinicalTrials.gov record from January 5, 2017.

#### Timing for Results Becoming Due

The Final Rule states that, for applicable trials, results information “must be submitted no later than 1 year after the primary completion date.”^9^ ClinicalTrials.gov requires a trial to have an expected completion date during trial registration. This field is then to be updated within 30 days of reaching the final or “actual” primary completion date. Whether a primary completion date is entered as “expected” or “actual” is included in the XML record but has no impact on whether a trial is considered due to report under the law. The FDAAA 2007 specifies that results are due one year from the earlier of the estimated or actual primary completion date.^8^ This is likely to prevent any loopholes regarding out-of-date or neglected registry entries never becoming due.

All submitted results are subject to quality control (QC) by ClinicalTrials.gov staff to ensure they meet a minimum standard. The authors of the Final Rule make clear that results information is supposed to be posted to ClinicalTrials.gov within 30 days following their submission, regardless of QC status. Sponsors may also, in certain instances, apply for certificates that delay the reporting of results. It was necessary to account for these delays when building our tracker. The final logic used to identify when a trial’s results are due is summarized in Table 3 followed by our methods to account for any issues that arose.

**Table 3:** Logic for Due Trials

| Category | Logic |
| --- | --- |
| **Due to Report Results** | The current date is later than the primary completion date + 1 year + 30 days<sup>a,b</sup><br>**AND**<br>Trial is an ACT **OR** pACT<br>**AND**<br>Trial does not have a disposition to delay results **OR** it has been 3 years + 30 days since primary completion date<sup>a,b</sup> |

<sup>a</sup> For all date values, when only a Month/Year were given, dates were defaulted to the last day of the given month (e.g. January 2017 = January 31, 2017).
<sup>b</sup> "Completion Date" field was used when "Primary Completion Date" was not available.

#### 30 Days Delay

Correspondence in 2017 with ClinicalTrials.gov indicated that the requirement to post results within 30 days, regardless of QC status, has not yet been implemented (Appendix 1). Previously, the ClinicalTrials.gov Final Rule website stated that: “More information on the remaining steps to implement fully the quality control review criteria and process, including posting of clinical trial information that has not yet met QC criteria, will be available soon”^15^. As of September 2019, ClinicalTrials.gov has announced that they plan to fully implement this requirement for all new results submitted from January 2020.^26^

Regardless of the implementation timeline for this requirement, we have kept the 30 day limit in our criteria for determining when results are due. This helps ensure accuracy on the tracker by allowing for a reasonable delay in processing and posting of trial information by ClinicalTrials.gov. 30 days also represents the timeline for notification of missing results before fines can be levied. Assuming prompt notification of responsible parties about missing results, a 30 day buffer allows for confidence in assessing when a trial is overdue to report and therefore eligible to be fined. In our experience running the tracker, delays of more than a few working days for information to be updated on ClinicalTrials.gov are rare.

We plan to follow closely how ClinicalTrials.gov implements the posting of results within 30 days regardless of QC status. Until this is confirmed, implemented, and adapted into our processes we will continue to rely on the “Results Submitted” tab on the trial record that details the QC process.^27^

The “Results Submitted” tab was added to ClinicalTrials.gov in 2017 to “help users track the submission and QC review status of results information.”^15^ Previously, this data was not available as part of the downloadable XML data record but a new “pending results” section was added to the XML to track trials undergoing QC as of 11 May 2018.^28^ This new field contains all dates related to the results submission QC process and is removed from the trial record and XML once the final results are posted. We have chosen to continue webscraping the “Results Submitted” tab for due trials, in favor of simply extracting data from the “pending results” XML, in order to ensure this data is properly archived.

Applicable trials that have full results posted prior to becoming due are immediately shown as “reported” on the tracker and counted in the reporting statistics regardless of when they become due. Day overdue is derived from the “results_first_submitted” field when full results are available. When the data is updated, each overdue trial without full results available, including trials that would become overdue that day, are checked for the presence of the “Results Submitted” tab. This allows us to identify these trials in QC as “reported.” The dates provided in the “Results Submitted” tab allow accurate tabulation of the “days overdue” field displayed next to each trial in the tracker even when the trial is in QC. This field will remain empty on the tracker until any results are submitted, or the 30 day administrative delay has passed and we are confident the trial is overdue. If the trial has not reported within the 30-day window, the 30-day administrative delay will then be accounted for in the “days overdue” calculation and adjusted based on any submission dates provided. Trials that do report within the 30 day window are still displayed as “reported late” with the appropriate days late provided relative to the primary completion date.

#### Delaying the Submission of Results

The Final Rule brought much needed clarity on reporting requirements for trials of unapproved drugs and devices and how this related to requesting certificates of delay. Sponsors of trials of unapproved products that are seeking, or plan to seek, an initial approval, licensure or clearance, or a new indication for an existing product from the FDA may request a certificate that delays the deadline to report so long as they are also the manufacturer of that product.^9^ If the certificate is granted, results become due at the earliest of: three years after the primary completion date; 30 days after a drug or device receives an FDA approval; or a marketing application/premarket notification is withdrawn without resubmission for at least 210 days. Sponsors may also apply for deadline extensions if they can demonstrate “good cause” although this is not distinguishable in the study record from a certificate of delay.

Any delay to results reporting attributable to this process is recorded in the “disposition” data field in the public XML and included in our data extraction. As the exact length or type of “disposition” is not available, and we do not account for the FDA approval/application status of products studied in trials, we assume the delay will last for three years from the primary completion date or until results are otherwise provided. It would be helpful if ClinicalTrials.gov gave more detail on the disposition duration in the downloadable and/or publicly accessible data for trials with such extensions.

#### Unclear Dates

Many records on ClinicalTrials.gov provide key dates only in month/year format without specifying a day. We have established in correspondence with ClinicalTrials.gov (Appendix 1) that sponsors are required to give a day, month, and year within 30 days of when they have an actual “Primary Completion Date”; sponsors who fail to do so are therefore breaching their obligation to post accurate data onto the public record. However it is common for sponsors to give incomplete data for completion dates. In these instances we defaulted their date to the last day of the given month (e.g. January 2017 = January 31, 2017). This allows a conservative assessment of when a trial started, ended, and when it is due to report results. It does present a minor issue for the small number of trials beginning or ending in January 2017 that fail to give complete date data: trials that actually started just prior to January 18, 2017 should be held to the pACT standard but will instead be held to the ACT standard, and pACTs that actually ended just prior to the effective date will be held to the standard of the Final Rule for reporting results. This decision may lead to a very small number of “January 2017” trials being incorrectly included or excluded from our tracker as a result of incomplete information provided on ClinicalTrials.gov by the trial sponsor. We expect this aspect of sponsors’ incomplete data will have negligible impact on the tracker overall, and any issues should improve over time, as most sponsors will hopefully update their records with accurate and precise start and completion dates.

#### Calculating Fines

While ClinicalTrials.gov is maintained by the National Institutes of Health, the FDA is tasked with carrying out enforcement actions related to clinical trial information, including non-submission of results.^29^ The FDA may assess an initial fine of “not more than $10,000 for all violations adjudicated in a single proceeding” for any missing information. Additional fines of up to $10,000 for each day that required trial information is not submitted, following a 30 day notification period, may also be assessed.^9^ In 2017, the fine amount for missing trial results was inflation-adjusted upwards from $10,000 to $11,569 per Department of Health and Human Services rulemaking.^30^ For implementation of these fines in the TrialsTracker, we only track the potential ongoing daily fines for trials that fail to report after 30 days. We believe there is considerable variation in the way FDA could administer the initial fine “in a single proceeding” which would make it difficult to automatically account for on the tracker. In order to remain conservative, we have decided not to include this initial fine in our calculations.

When sponsors submit results, exact submission dates are available as a data element from ClinicalTrials.gov, either in the XML record (when results have been posted) or via the “Results Submitted” tab (when results are in QC). As such, after 30 days from the 1 year deadline we calculate a potential fine of $11,569 for each day with no indication that results have been submitted. This assumes an immediate notification of the sponsor that the deadline for results submission has been missed which should be possible via the PRS sponsor accounts. We will also monitor the FDA website for any indication that fines have been levied and provide this information on the tracker, in order to place potential fines in the context of actual fines levied. There is no indication that any fines have been levied as of October 2019.

#### Canceled Results

In the 11 May 2018 update to ClinicalTrials.gov, it was announced that canceled results will now be recorded in the “Results Submitted” tab.^28^ A canceled results submission is when a sponsor or investigator recalls submitted results before the QC process can take place. In order to account for this on the TrialsTracker website we have created a new category of overdue trials which indicate that previously submitted results have been canceled. It is unclear how or when these trials would be eligible for fines so we conservatively do not calculate any new additional fines for a trial with canceled results beyond those that may have initially accrued before their first submission. The “days overdue” field will update as if results were never submitted (regardless of which QC round the cancellation took place in) until a new, non-canceled results submission occurs. When full results become available, we revert to assessing the on-time status of the trial based on its initials submission date. As the status of canceled results, and how they interact with the required submission timelines, is not well documented, we believe this process accurately reflects our current understanding of the process and ensures that sponsors are both properly credited for submitting results on-time and held accountable to make their results public as quickly and efficiently as possible.

### Website

Using data from ClinicalTrials.gov and our derived values for coverage and reporting, we created a live tracking website, The FDAAA TrialsTracker (fdaaa.TrialsTracker.net), to provide up-to-date statistics on what sponsors are not reporting the results of “due” trials on ClinicalTrials.gov. The website launched initially on February 19, 2018 in a Beta period, with a full launch on 5 March 2018. Updates are each working day, with future update frequency to be determined by available resources. All ACTs and relevant pACTs identified are included on the website. Users are able to view summary statistics, all individual trials, and trials categorized by sponsor; and download data for their own use either through the website GUI or an included API. Filters are available for a variety of trial statuses. The total possible fines that could have been collected overall and from each individual sponsor are also displayed. Full ClinicalTrials.gov data downloaded for each update is archived for potential future analysis and can be shared on request. Figures 1 and 2 include screenshots of the “Ranked Sponsors” and “Single Trials” views.

**Figure 1:**
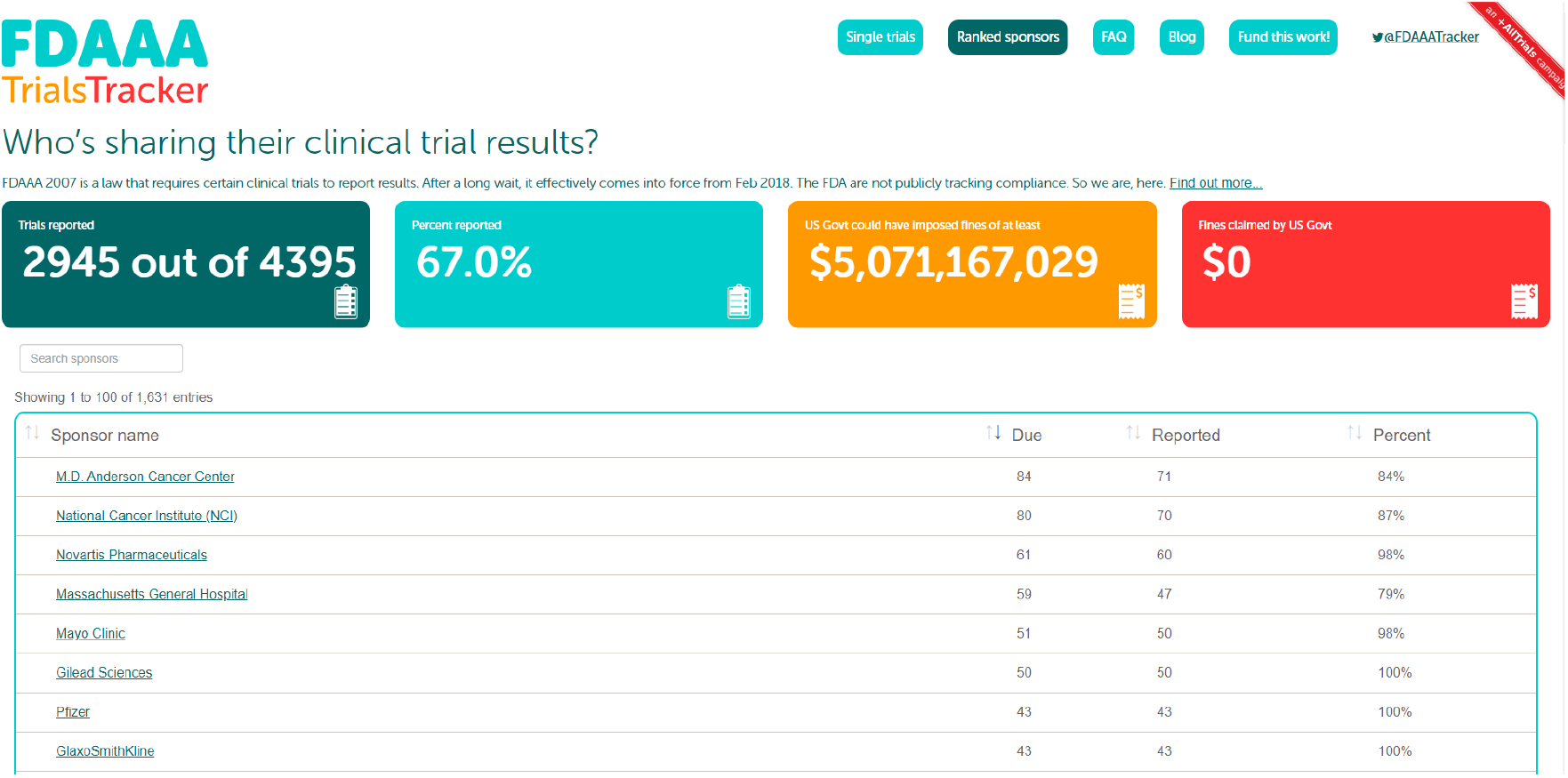
Ranked Sponsor Page (Data as of 1 Oct 2019)

**Figure 2:**
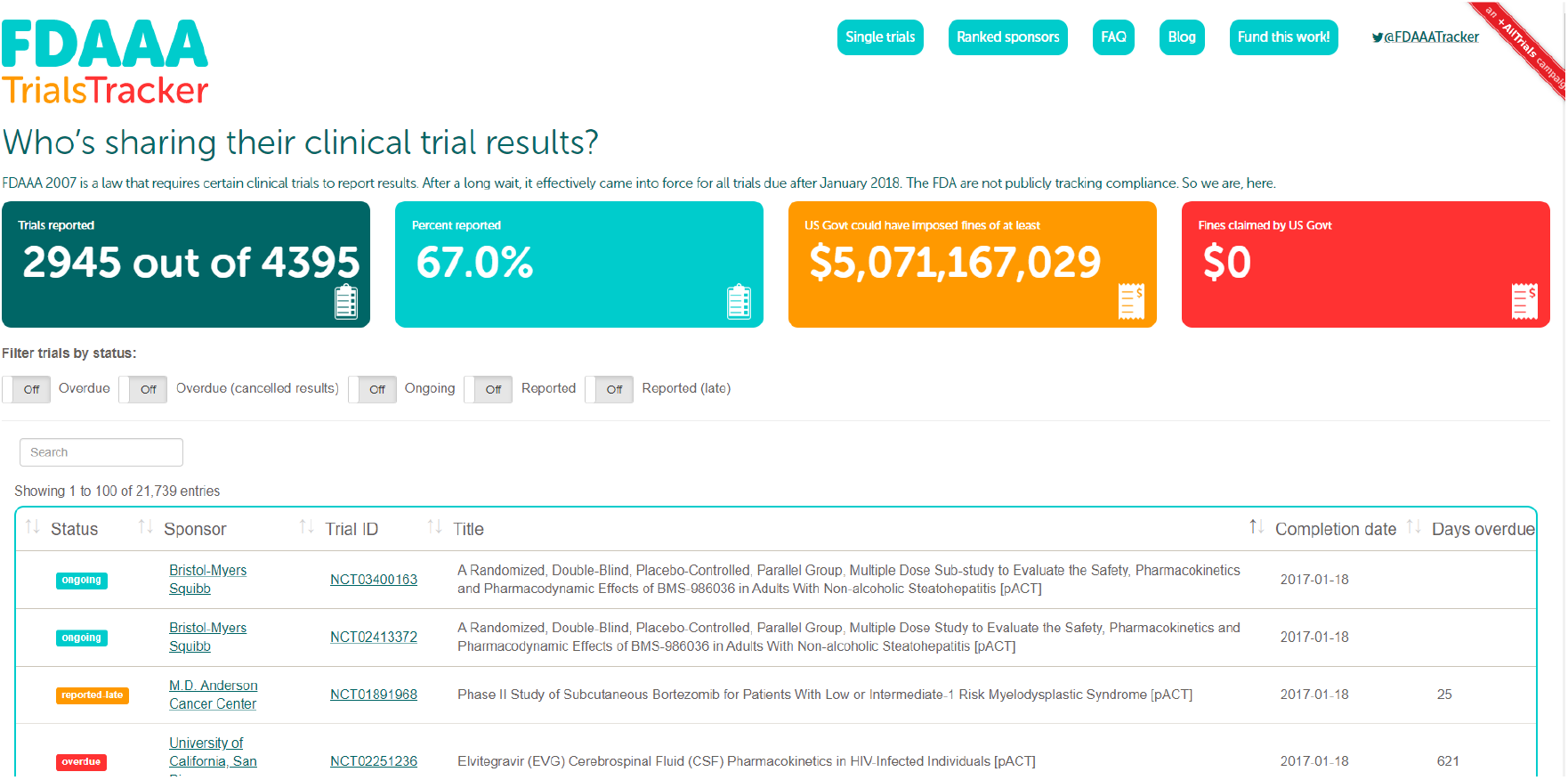
Single Trials Page (Data as of 1 Oct 2019)

### Unresolved Issues

While the global ethical standard is that all trials should report results^1^, we have set out to implement all the reporting exceptions of FDAAA. Following the decision by ClinicalTrials.gov and/or FDA to withhold information on ACT and pACT status from the public record, there remain a very small number of outstanding trials where FDAAA coverage cannot be perfectly ascertained from publicly available structured data. We document any specific outstanding issues below.

#### Bioequivalence

Phase 1 trials are universally excluded from the reporting requirements of the FDAAA 2007; certain bioequivalence trials are done after marketing approval, and share characteristics with phase 1 trials, but may not labeled as such in the data. The Final Rule provides some guidance on this issue, noting that “bioequivalence or comparative bioavailability” studies that meet a certain definition are excluded from reporting while those meeting a different definition are not excluded.^9^ To our knowledge, this distinction cannot be ascertained via any field in the publicly shared data. The lack of clarity and actionable data fields related to this distinction could rarely lead to trials being misidentified on the tracker as due to report under FDAAA 2007 when they are actually exempt. We are documenting and monitoring this issue.

## Discussion

### Summary

Following extensive review of the legislation it was possible to develop and deliver a live website which: publicly audits compliance with the results reporting requirements of the FDAAA 2007 and the Final Rule; implements the FDAAA reporting exceptions; identifies individual trials which are overdue; presents sponsor-level performance; and updates automatically.

### Strengths and Weaknesses

Our tool and website openly tracks compliance with transparency reporting legislation across all trials, with live updates as the data changes. We cover all publicly identifiable trials conducted under FDAAA 2007 on ClinicalTrials.gov, and our data updates each working day. We faced challenges in the form of ClinicalTrials.gov withholding data and sponsors entering poor quality and incomplete data onto the register. We used a conservative approach to work around these issues and some sponsors’ giving incomplete data on dates; for the reasons given above, we think our assumptions were reasonable and conservative, in that they minimize the chances of us incorrectly identifying a trial as due to report results; furthermore, we believe these issue affect only a small number of trials, and will therefore have only a negligible impact on the tool.

A key strength of our methods was our collaborative approach. The DataLab is a multidisciplinary team consisting of academics, clinicians, and software engineers working together to produce live interactive tools from data, as well as static analyses for academic publications, across a range of medical problems including health informatics as well as trials transparency.^31^ The analysis, tool, and website reported here were developed and delivered internally and iteratively, rather than through external procurement. This improves efficiency and builds capacity to deliver further innovative tools, as we have a team that consists: of software engineers who understand aspects of evidence based medicine; and researchers who understand aspects of delivering data-driven websites. Further analyses based on insights from the TrialsTracker project is currently being prepared for publication.

### Context of Other Work

To our knowledge this is the first tool and website to openly and publicly track compliance with transparency legislation across all trials, with live updates as the data changes. Previous work assessing compliance with FDAAA 2007 was produced prior to the final rule, and delivered only static analyses for the purpose of one-off academic publications, with data that has rapidly gone out of date.^32,33^ Previous work on publication bias has generally relied on laborious manual searches to assess reporting, and has consequently run on a limited sample of trials, and again on a one-off or very infrequent basis.^4,21,22,34^ Our tool runs on all trials on ClinicalTrials.gov and updates each working day. Prior work also underscores the importance of reporting to results to ClinicalTrials.gov. A number of studies have found that results to ClinicalTrials.gov are often more complete than corresponding journal publications, especially regarding the reporting of adverse events.^35–37^

We have previously produced an automated and updatable tool that estimates the proportion of trials that have reported results across a very large sample of trials, by searching for results of completed trials on clinicalTrials.gov itself, and also by searching for those trials’ results in academic papers, using a series of automated and filtered searches on PubMed. This tool deliberately casts its net more widely than the narrow requirements of FDAAA 2007, mirroring the ethical obligations to report all trials, and therefore checks whether all trials since 2006 have reported their results. As reported in that previous manuscript, the approach used in that tool reflects a trade off between covering a very large number of trials, in a regularly updating service, at the cost of lower accuracy than manual search; whereas manual search can cover only a small number of trials, and cannot be regularly updated to produce ongoing public audit.^38^ However, under FDAAA 2007, trials are required to report their results directly onto ClinicalTrials.gov itself; compliance with the requirement to report results can therefore be ascertained unambiguously and completely.

### Policy Implications

Past work has shown that results from trials often go unreported^4^, despite numerous guidelines, commitments and legal frameworks intended to ensure complete reporting. Without formal sanctions being imposed by the FDA and others, we believe that open data tools that provide public accountability have a valuable role in improving standards^11^. Specifically, we hope that the presence of easily accessible public data, and rankings, showing how individual sponsors are meeting their obligations, may encourage organizations to prioritize results reporting in general. In particular, the dynamic nature of the data presented through our tools incentivizes organizations to report their trial results, because - unlike in a static academic publication on trial reporting - they can immediately improve their public rating, by reporting their results. In addition, the online resources we have produced here and elsewhere make it extremely easy for sponsors to identify individual trials from their organizations which have not yet reported their results: our tools therefore offer positive practical support for those sponsors who wish to ensure that all their completed trials have reported results.

We continue to build the TrialsTracker programme with additional trackers. In September 2018, we launched the EU TrialsTracker (eu.trialstracker.net) which tracks compliance with EU Guidelines requiring sponsors to report results for European trials of medicinal products directly to the EU Clinical Trial Register. Like the FDAAA TrialsTracker, summary reporting statistics, sponsor rankings, and trial-level reporting is presented to users. Data is refreshed monthly.^39^ We are keen to receive feedback to improve all such tools. We are keen to hear feedback on all of our TrialsTracker projects from trialists, institutions, funders, regulators, patients, the public, and others.

### Future Research

We plan to seek publication of these methods in a peer-reviewed journal which will also include an analysis of compliance rates and other facets of the legislation.

### Conclusions

Open data tools that provide live data on trials transparency can improve accountability, and have great potential to help ensure that all trials are reported.

## Article Information

## Acknowledgements

We are grateful to Holly Fernandez-Lynch of University of Pennsylvania for advice on interpretation of FDAAA 2007; Francis Irving developed our EUCTR tracker, the codebase of which is adapted for our FDAAA tracker, and contributed to discussions on websites to drive accountability; Helen Curtis, Alex Walker, Richard Croker, Emma-Jane Greig, Peter Ingle-sham, Brian McKenna at EBM DataLab contributed feedback on the website.

## Competing Interests

All authors have completed the ICMJE uniform disclosure form at www.icmje.org/coi_disclosure.pdf and declare the following: BG has received research funding from the Laura and John Arnold Foundation, the Wellcome Trust, the Oxford Biomedical Research Centre, the NHS National Institute for Health Research School of Primary Care Research, the Health Foundation, and the World Health Organisation; he also receives personal income from speaking and writing for lay audiences on the misuse of science. NJD and SB are employed on BG’s grants from LJAF for this work. SB is also employed under BG’s grants related to the OpenPrescribing project. NJD has previously been employed on grants from the Open Society Foundation and the State Attorney General Consumer and Prescriber Education Grant Program.

## Funding

BG is funded by the Laura and John Arnold Foundation to conduct work on research integrity. No specific funding was sought for this project. The funder had no role in the study design, collection, analysis, and interpretation of data; in the writing of the report; and in the decision to submit the article for publication.

## Ethical approval

This study uses exclusively open, publicly available data, therefore no ethical approval was required.

## Contributorship

BG conceived the project. BG SB ND devised the methods and collected the data. ND conducted the policy review. ND developed a preliminary off-line analysis of the data with input from SB and BG. SB built the website with input from ND and BG. SB maintains the codebase and ND contributes code to the project. ND and BG drafted the manuscript. All authors contributed to and approved the final manuscript. BG supervised the project and is guarantor.

## Appendix 1

### Correspondence with ClinicalTrials.gov Support

**Ticket #28045-279395**

**11 Nov 2017**

Dear ClinicalTrials.gov Staff,

I am interested in assessing some characteristics of applicable clinical trials (ACT) per 42 CFR 11.22(b) since the effective date of January 18, 2017.

I was able to locate the published checklist here (https://prsinfo.clinicaltrials.gov/ACT_Checklist.pdf) but a number of the data elements used to determine whether any given record is an ACT are unvailible in the public XML.

Using the advanced search, I created a full XML record of all phase 2-4 interventional studies posted from January 18, 2017 until the end of October 2017. This covered 5,640 records in total. None of the publicly available XML contained the following data fields referenced in the above checklist:

“U.S. Food and Drug Administration IND or IDE Number” “Product Manufactured in and Exported from the U.S.” “Studies a U.S. FDA-regulated Device Product” “Studies a U.S. FDA-regulated Drug Product”

I was also unable to locate any specific flag or field that would note if a given record meets the criteria of an ACT. Are there plans to create such a flag, or make the required elements necessary to determine an ACT public, so trials can be easily identified for analysis? It appears that it is currently impossible for a member of the public to definitively identify an ACT in ClinicalTrials.gov given the available public information.

Best,

Nicholas DeVito

**11 Nov 2017**

Hi Nick,

Yes, you are correct, we do not have some of the data fields available.

Hopefully this will be corrected in the future.

ClinicalTrials.gov

**Ticket #28045-288723**

**1 Dec 2017**

Per my previous question (Ticket #28045-279395) I would like to follow-up.

Our goal is to determine whether a certain trial is an applicable clinical trial (ACT) as this information is important for ascertaining whether researchers are meeting their statutory obligation to report results within 12 months.

Can you please clarify the following:

1. Do you know internally whether a given trial is an ACT? If so, is this obtained by utilizing the existing data fields as outlined here (https://prsinfo.clinicaltrials.gov/ACT_Checklist.pdf) or in some other manner?
2. To confirm, based on the information available to the public on ClinicalTrials.gov there is currently no definitive way to establish whether a given trial is an ACT?
3. Would we be able to apply or petition for an ACT flag, or the appropriate underlying data fields, to be made public in some way?

Thank You,

Nicholas DeVito

**5 Dec 2007**

Hi there,

1. All trials internally are marked ACT, PACT or NON ACT. We do this by using the check list. The administrator at your organization have this information and we supply reports to them.
2. Yes, this is correct.
3. I will pass this on to our systems team, however in some case if we did this, proprietary information would be exposed.

ClinicalTrials.gov

**Ticket #28045-292644**

**12 Dec 2017**

Hello,

I was curious as what the delay is for posting results to clinical trials.gov after they are received from the responsible party? Is this defined in law? What would be a safe amount of time to add to the 1 year statuary requirement as an administrative buffer for results to be posted?

Best,

Nicholas DeVito

**11 Nov 2017**

Hello,

Please see information in the FAQs at: https://clinicaltrials.gov/ct2/manage-recs/faq#resultsInfoSubDeadline

Thank you,

ClinicalTrials.gov

**Ticket #28045-293891**

**14 Dec 2017**

Hello,

Pursuant to my previous ticket #28045-292644, I would like to request further clarification concerning posting results information beyond what is available in the FAQ.

My team will shortly be launching a tool which tracks and identifies trials that appear to have breached the FDAAA2007 requirement to post results to clinicaltrials.gov within 12 months of trial completion as described in 42 CFR Part 11. We have read the FAQ as well as the relevant sections of the FDAAA 2007 final rule (specifically those pertaining to section 5.§11.52).

To confirm, if an Applicable Clinical Trial with no Certificate of Delay (or other noted dispensation) and no results posted publicly on ClinicalTrials.gov after 12 months plus 30 calendar days after its primary completion date, is it reasonable to assume it has breached the FDAAA requirement to post results? Or could there be further delays before a trial’s results appear on clinicaltrials.gov that we should be aware of?

Thank You,

Nicholas DeVito

**15 Dec 2017**

Hi there,

Please note, they could have submitted the data to us, however because the of the review process, it may take more than 30 days.

ClinicalTrials.gov

**Ticket #28045-295627**

**19 Dec 2017**

Hello,

My prior inquiries #28045-293891 and #28045-292644 are related to the timeline for posting results on ClincialTrials.gov following their submission be the responsible parties. The last response noted that:

“They [the responsible party] could have submitted the data to us, however because the of the review process, it may take more than 30 days.”

However the FDAAA Final Rule strongly states that the results information will be posted online within 30 days of the due date, with no further delays for quality control, and indeed discusses the benefits and hazards of posting results before they have had a more lengthy review. I have posted the relevant sections of the Rule below. Can you please tell me if there is some additional cause for delay that we are unaware of, that is not covered by this aspect of the Final Rule? Or, if something has been changed, could you tell us what the new deadline is, and where we can read more about how this aspect of the Final Rule has been revised?

§11.52 - By when will the NIH Director post submitted clinical trial results information? Overview of Statutory Provisions and Proposal According to section 402(j)(3)(G) of the PHS Act, for applicable clinical trials, the Director of NIH is required to post results information “publicly in the registry and results database not later than 30 days after such submission.”

Commenters expressed concern about the potential to misinform those using the public record and suggested only posting sections that have fulfilled quality control criteria. Some commenters suggested that the harm of posting information before the quality control review process has concluded is greater than the benefit of posting the information in a timely manner. While we understand these concerns, we interpret the statutory posting deadline to be a clearly delineated timeline between submission and posting. In addition, in the event that a study record is posted in accordance with the statutory posting deadline and the quality control review process has not concluded, the clinical trial record will contain information that will be visible to those viewing the record on ClinicalTrials.gov to make it clear that the quality control review process has not concluded for the posted clinical trial information.

Many thanks,

Nicholas DeVito

**No Response from ClinicalTrials.gov**

**Ticket #28045-301558**

**8 Jan 2018**

Hello,

I have previously been in touch concerning details related to the new results reporting requirements on ClinicalTrials.gov. My previous inquiries are #28045-292644, #28045-293891, and #28045-295627.

The last of these (#28045-295627) has not yet been replied to however I understand that this may have gotten lost in the bustle of the holidays. I have repeated this question along with two additional inquiries below:

1. The last response to one of my inquiries regarding the timeline for reporting results (#28045-293891) noted that:

“They [the responsible party] could have submitted the data to us, however because the of the review process, it may take more than 30 days.”

However the FDAAA Final Rule strongly states that the results information will be posted online within 30 days of the due date, with no further delays for quality control, and indeed discusses the benefits and hazards of posting results before they have had a more lengthy review.

§11.52 of the Final Rule states that: “The Director will post publicly on ClinicalTrials.gov the clinical trial registration information, except for certain administrative data, for an applicable drug clinical trial not later than 30 calendar days after the responsible party has submitted such information, as specified in §11.24.”

Earlier in the same document, the rationale and interpretation of this requirement is described at length:

“Commenters expressed concern about the potential to misinform those using the public record and suggested only posting sections that have fulfilled quality control criteria. Some commenters suggested that the harm of posting information before the quality control review process has concluded is greater than the benefit of posting the information in a timely manner. While we understand these concerns, we interpret the statutory posting deadline to be a clearly delineated timeline between submission and posting. In addition, in the event that a study record is posted in accordance with the statutory posting deadline and the quality control review process has not concluded, the clinical trial record will contain information that will be visible to those viewing the record on ClinicalTrials.gov to make it clear that the quality control review process has not concluded for the posted clinical trial information.”

Can you please tell me if there is some additional cause for delay that we are unaware of, that is not covered by this aspect of the Final Rule? We noticed the recent posting about new features on ClinicalTrials.gov (https://www.nlm.nih.gov/pubs/techbull/nd17/nd17_clinicaltrials_enhanced.html) included a section on the new “Results Submitted” tab. This would appear to contradict the Final Rule and allow for quality control to delay the posting of results longer than 30 days.

2. Regarding the new “Results Submitted” feature, we noticed that this does not appear to currently be represented in the XML of study records. Specifically, XML records for studies that include this new tab say “No Results Available” for the <study_results> section with no other indication in the record that results have been submitted but are currently undergoing quality control (ex: NCT01798225).

Is this correct? Are there any plans to add notation to the XML describing the information currently represented on the “Results Submitted” tab? If so when would that be expected?

3. Regarding the checklist for ACTs (https://prsinfo.clinicaltrials.gov/ACT_Checklist.pdf) can you confirm that when responsible parties are inputting trial data to ClinicalTrials.gov, they must have at least one aspect of criteria 2 checked (facility in US, IND/IDE, manufactured/exported from US) in order to be able to provide an affirmative response to criteria 3 (regarding FDA regulation of a drug or device product)?

Thank you in advance for your help regarding these matters.

Best,

Nicholas DeVito

**11 Jan 2018**

Answers to your questions:

1. The 30-day posting requirement has not yet been implemented. Please see the PRS Info Page (https://prsinfo.clinicaltrials.gov/) for updates on Final Rule implementation. Note the following from this page:

“Quality control (QC) review criteria and process (42 CFR 11.64(b))

- April 18, 2017: Study record review comments provided by the National Library of Medicine (NLM) as part of the QC review process are labeled as either Major or Advisory comments when returned to the responsible party. While each major issue identified in the comments must be corrected or addressed, advisory issues are suggestions to help improve the clarity of the record.
- December 18, 2017: Study records with results submitted but not yet posted on ClinicalTrials.gov include a Results Submitted tab (in place of the No Results Posted tab) to help users track the submission and QC review status of results information. The tab displays a table of dates showing when results information was submitted and, if applicable, returned to the responsible party with QC review comments identifying at least one major issue. In addition, the following dates are summarized on the Key Record Dates page for each record:

First Submitted that Met QC Criteria
Results First Submitted that Met QC Criteria
Last Update Submitted that Met QC Criteria

For more information see ClinicalTrials.gov: Further Enhancements to Functionality.
- More information on the remaining steps to implement fully the quality control review criteria and process, including posting of clinical trial information that has not yet met QC criteria, will be available soon?

2. You are correct, this is not available in xml.

3. Required and optional data elements are described in the ClinicalTrials.gov Protocol Registration Data Element Definitions for Interventional and Observational Studies (https://prsinfo.clinicaltrials.gov/definitions.html).

The “Product Manufactured in and Exported from the U.S.” is required if U.S. FDA-regulated Drug and/or U.S. FDA-regulated Device is “Yes,” U.S. FDA IND or IDE is “No”, and Facility Information does not include at least one U.S. location.

Please see the FDAAA 801 Problems section of the PRS User’s Guide (at: https://prsinfo.clinicaltrials.gov/prs-users-guide.html#fdaaa801problems) for full explanation on the data elements used to identify probable applicable clinical trials (pACTs) and applicable clinical trials (ACTs) in the PRS.

**Ticket #28045-304148**

**15 Jan 2018**

Hello,

Thank you for your response to my previous enquiry (#28045-301558).

One of my questions in that enquiry read:

“Regarding the checklist for ACTs (https://prsinfo.clinicaltrials.gov/ACT_Checklist.pdf) can you confirm that when responsible parties are inputting trial data to ClinicalTrials.gov, they must have at least one aspect of criteria 2 checked (facility in US, IND/IDE, manufactured/exported from US) in order to be able to provide an affirmative response to criteria 3 (regarding FDA regulation of a drug or device product)?”

To which I received the response:

*“Required and optional data elements are described in the ClinicalTrials.gov Protocol Registration Data Element Definitions for Interventional and Observational Studies* (https://prsinfo.clinicaltrials.gov/definitions.html).

*The “Product Manufactured in and Exported from the U.S.” is required if U.S. FDA-regulated Drug and/or U.S. FDA-regulated Device is “Yes,” U.S. FDA IND or IDE is “No”, and Facility Information does not include at least one U.S. location*.

*Please see the FDAAA 801 Problems section of the PRS User’s Guide (at*: https://prsinfo.clinicaltrials.gov/prs-users-guide.html#fdaaa801problems) *for full explanation on the data elements used to identify probable applicable clinical trials (pACTs) and applicable clinical trials (ACTs) in the PRS*.”

We had previously reviewed the “Protocol Registration Data Element Definitions” and understand what is and is not required by the responsible parties entering data. However, this response does not fully answer our question.

To clarify, we would like to know if, functionally, when a responsible party is entering information into the ClinicalTrials.gov website, would they be able to enter information into the “FDA-regulated Drug and/or Device” field without first meeting one of the conditions of criteria 2 (facility in US, IND/IDE status, manufactured/exported from US)?

We ask because we are interested in being able to identify ACTs using the public data, however since “U.S. FDA IND or IDE” data element is not public, it would not be possible to definitively identify an ACT. However, in discussions with colleagues, we have heard that the criteria in question 2, while required, may be redundant to criteria 3 for publicly determining ACT status since criteria 3 cannot be entered without first meeting one of the requirements outlined in criteria 2. We would like confirmation of this fact as it would be helpful our ACT identification protocol.

Thank You,

Nicholas DeVito

**18 Jan 2018**

If you enter no US locations, and answered NO to the question Product Exported from U.S and you answered YES to either U.S. FDA-regulated Drug or U.S. FDA-regulated Device, then you would get the following error.

ERROR: U.S. FDA-regulated Drug cannot be ‘Yes’ unless this study is an IND study, has one or more U.S. Locations, or is a study of a drug that is exported from the U.S.

ClinicalTrials.gov

**Ticket #28045-324772**

**8 Mar 2018**

Hello,

Are responsible parties required to add a date to their Primary Completion Date field on a ClinicalTrials.gov registry entry once that date has been reached or is it allowable to remain in Month/Year format?

For instance, if a trial had a primary completion date of “January 2017” are they technically violating their responsibility to maintain their record if they have not yet specified which day in January 2017 or updated their entry with additional information concerning a new primary completion date?

If responsible parties are not required to provide a “Day” in this field, how are the various deadlines dependent on the primary completion date calculated?

Many thanks,

Nicholas DeVito

**8 Mar 2018**

The following is listed in the protocol registration data elements. http://clinprsqa/prs/html/definitions.html

Once the clinical study has reached the primary completion date, the responsible party must update the Primary Completion Date to reflect the actual primary completion date.

ClinicalTrials.gov

**Ticket #28045-325151**

**9 Mar 2018**

Hello,

Per my previous ticket #28045-324772, can you please confirm that a specific date is required to be entered for the “primary completion date” and “completion date” fields by sponsors on clinicaltrials.gov, when they become available?

I am familiar with the definition provided here: http://clinprsqa/prs/html/definitions.html

However, this is not explicit that an “Actual Primary Completion Date” is expected to include the exact day of completion and not just the month/year.

Thank you,

Nicholas DeVito

**9 Mar 2018**

Yes, the exact date is required.

ClinicalTrials.gov

**Ticket #CAS-313291-S6Q3J7**

**14 Jan 2019**

Hello,

I have previously written and was informed that, per ACT criteria, a trial that selects yes to the “FDA Regulated Drug Product/Device” field would necessarily mean that one of the “exported from the US,” “part of an IND/IDE,” or “study site in the US” criteria would be true. If a trial that meets pACT criteria updates to include the new “FDA Regulated Drug/Device” fields, does it make more sense to switch the assessment of these trials to use the ACT criteria than the pACT criteria, assuming the above assumption holds true? It seems to me that when these new fields are available and updated by the sponsor, that would mean the ACT criteria can be applied even for trials that began prior to January 18, 2017.

Thank you,

Nicholas DeVito

**16 Jan 2019**

Hi there, It is only an ACT if it started on or after 1/18/2017.

**17 Jan 2019**

Hi,

Yes, I understand this per my original message. My question is, since certain information (specifically IND/IDE status) is not available to the public, can we rely on the newly added fields (i.e. FDA Regulated Drug/Device) when they are added to a pre-Jan 18, 2017 trial, in order to determine if a trial that meets all the other pACT criteria, but say, has no US location, to determine whether that trial is a pACT (or more clearly, whether that trial is required to report results under the Final Rule).

Best,

Nicholas DeVito

**18 Jan 2019**

Back to your original email, the “pACT” and “ACT” labels are just naming conventions used only in the PRS to assist responsible parties in identifying trials subject to section 402(j) of the PHS Act (pACT) and the newer requirements in 42 CFR Part 11 (ACT).

The information provided in this FAQ seems relevant: https://clinicaltrials.gov/ct2/managerecs/faq#fr_1_

“Beyond their primary purpose, the ACT Checklist and Elaboration may also be useful to assist in evaluating whether a clinical trial or study that was initiated before January 18, 2017, and which is not subject to the final rule requirements, is an ACT under section 402(j) of the Public Health Service Act.

We note that Responsible Parties or other users of the ACT Checklist and Elaboration are responsible for using accurate data about a clinical trial or study and for properly evaluating whether the trial or study must be registered and, if so, which results must be submitted.”

## Notes

#### Summary of Updates

This document has been updated to reflect changes to the FDAAA TrialsTracker implemented since the last revisions in response to new information and changed form ClinicalTrials.gov.

https://fdaaa.trialstracker.com

https://github.com/ebmdatalab/clinicaltrials-act-tracker

https://github.com/ebmdatalab/clinicaltrials-act-converter

